# MrgD Receptor Modulates Neurotransmission in the Nigrostriatal Pathway

**DOI:** 10.1101/2025.07.10.664011

**Authors:** Lucas Rodrigues-Ribeiro, Bruna da Silva Oliveira, Kivia Soares Barretos Santos, Caroline Amaral Machado, Arkadiusz Nawrocki, Maria José Campagnole dos Santos, Aline Silva de Miranda, Cristina Guatimosim, Martin Røssel Larsen, Robson Augusto Souza dos Santos, Thiago Verano-Braga

**Affiliations:** National Institute of Science and Technology in Nano Biopharmaceutics, Department of Physiology and Biophysics, Federal University of Minas Gerais, Belo Horizonte, MG, Brazil; Department of Biochemistry and Molecular Biology, University of Southern Denmark (SDU), Odense, Denmark; Department of Morphology, Federal University of Minas Gerais, Belo Horizonte, MG, Brazil

**Keywords:** MrgD receptor, nigrostriatal pathway, neurotransmission, motor hyperactivity, compulsivity, β-alanine, alamandine

## Abstract

The Mas-related G protein-coupled receptor D (MrgD) is primarily known for its role in peripheral nociception and, more recently, as a receptor for alamandine, influencing cardiovascular function and exhibiting antidepressant-like effects. However, its function within the central nervous system, particularly in motor and reward-related circuits, remains largely unexplored. Here, we investigate the role of MrgD in the nigrostriatal pathway using proteomic approaches focused on post-translational modifications in MrgD-knockout (KO) mice. Integrated proteomic, phosphoproteomic, and N-glycoproteomic analyses revealed significant alterations in synaptic vesicle-associated proteins, pointing to impaired neurotransmission in the nigrostriatal system of KO mice. These molecular findings were supported by neurotransmitter quantification and functional assays demonstrating impaired synaptic vesicle exocytosis. Pharmacodynamic analyses showed that MrgD modulates synaptic exocytosis in an agonist-selective manner, being responsive to alamandine but not β-alanine. Furthermore, behavioral analyses revealed increased locomotor activity and compulsive-like behavior in MrgD-deficient mice, without impairments in short- or long-term memory. Together, these findings uncover a new role for MrgD beyond its involvement in nociception, highlighting this receptor as a potential therapeutic target for neurological disorders involving motor hyperactivity and compulsivity.

## INTRODUCTION

The Mas-related G protein-coupled receptor D (Mrgprd, also known as MrgD) is a member of the Mas-related G protein-coupled receptor (MRGPR) family, predominantly expressed in small-diameter sensory neurons of the dorsal root ganglia (DRG) [1]. It has been extensively studied for its role in nociception, mediating mechanical hyperalgesia and neuropathic pain [2, 3]. MrgD functions as a specific membrane receptor for β-alanine and is involved in the sensation of peripheral pain and itch [4]. Beyond its well-established role in sensory perception, recent evidence suggests broader physiological implications, including its involvement as a receptor for alamandine, impacting cardiovascular function and potentially exhibiting antidepressant-like effects [5, 6]. While its expression in the brain is not as robust as other MRGPR family members [7], MrgD has been localized in the cortex, hippocampus, amygdala, thalamus, hypothalamus and mid-brain areas, hinting at a potential, albeit less explored, role in central nervous system processes such as synaptic plasticity, learning, memory and locomotor modulation [8].

The nigrostriatal pathway, a crucial dopaminergic projection originating from the substantia nigra pars compacta (SNc) and innervating the striatum, is fundamental for motor control and coordination [9]. Dysfunction within this pathway is a hallmark of various neurological disorders, most notably Parkinson’s disease, characterized by the degeneration of dopaminergic neurons and subsequent motor deficits [10]. Neurotransmission within the nigrostriatal pathway relies heavily on the precise regulation of synaptic vesicle dynamics, including the synthesis, storage, release, and reuptake of neurotransmitters [11]. Synaptic vesicle proteins play a pivotal role in these processes, ensuring efficient and regulated communication between neurons [12]. Alterations in these proteins can lead to profound changes in synaptic function and, consequently, neurological pathology [10]. Monoamines, including dopamine, serotonin, and norepinephrine, are critical neurotransmitters that modulate a wide array of brain functions, with dopamine being the primary monoamine in the nigrostriatal pathway [13]. The intricate balance and interaction of these monoamines are essential for normal neurological function, and their dysregulation is implicated in numerous neuropsychiatric and neurodegenerative conditions [14].

Despite the established importance of MrgD in sensory systems and the critical role of the nigrostriatal pathway in motor control and disease, the role of MrgD in the nigrostriatal pathway has not been investigated. Our current study addresses this gap by presenting novel findings demonstrating that the MrgD receptor significantly impacts neurotransmission within the nigrostriatal pathway. Proteomics post-translational modifications analysis indicated an impairment in the neurotransmission in the nigrostriatal pathway. The lower synaptic vesicle exocytosis in the MrgD-knockout mice (KO) and the accumulation of dopamine (DA) in the substantia nigra and the decrease of DA in the striatum corroborated the molecular findings. Pharmacodynamic analysis showed that MrgD modulates synaptic exocytosis in an agonist-selective manner, being responsive to alamandine but not β-alanine. Furthermore, behavioral analyses revealed increased locomotor activity and compulsive-like behavior in MrgD-deficient mice, without impairments in short- or long-term memory. These results collectively demonstrate a new role for the MrgD receptor in the central nervous system, beyond nociception and opens new avenues for understanding the molecular regulation of dopaminergic signaling and for developing therapeutic strategies targeting disorders of the nigrostriatal pathway.

## METHODS

All reagents used in this study were purchased from Sigma-Aldrich, except for the phosphatase inhibitor (PhosSTOP™, Roche) and trypsin (produced by the proteomics research group (PR group), Odense, DK). Solutions were prepared using purified water from the Milli-Q system (Millipore Corporation, Bedford, MA). Low adhesion microtubes were used for sample processing in proteomic analyses.

### Ethical Aspects

The animal model experiments included in this study were conducted in accordance with ethical standards and regulations for the use of animals in research as described by the Ethics Committee on Animal Use of UFMG, under protocol number 241/2022.

### Animals

Male mice of the C57Bl/6 (WT) and MrgD-knockout (KO) strains were provided by the Biological Sciences Institute Animal Facility-2 (BICBIO-2) at UFMG. Adult male mice aged 3 and 12 months were used for the experiments. Genotyping was confirmed using the following PCR primers: MrgD-8, 5’-CATGAGATGCTCTATCCATTGGG-3’; reverse primer for tetracycline transactivator (rtTA1), 5’-GGAGAAACAGTCAAAGTGCG-3’; and MrgD-1, 5’-CTGCTCATAGTCAACATTTCTGC-3’. The animals had free access to water and food. Additionally, all mice were maintained at a controlled temperature of 23°C and a 12-12 hour light-dark cycle (the animal facility lights turn on at 7:00 AM and off at 7:00 PM).

### Tissue Collection

The animals were euthanized, and the brains were immediately collected and washed in an ice-cold phosphate-buffered saline (PBS) solution containing 137 mM NaCl, 2.7 mM KCl, 8 mM Na2HPO4, 2 mM KH2PO4, and protease and phosphatase inhibitors (pH = 7.4). The brain regions, including the substantia nigra (SN) and striatum (ST), were quickly dissected, frozen in liquid nitrogen, and stored in an ultrafreezer at −80°C.

### Proteomic Analysis Sample Preparation

Brain regions collected from mice of both groups (WT and KO) were homogenized in 300 μL of lysis buffer containing 1% (w/v) sodium deoxycholate (SDC), 20 mM tris(2-carboxyethyl) phosphine hydrochloride (TCEP), 40 mM chloroacetamide (CAA), 50 mM triethylammonium bicarbonate (TEAB, pH = 8), and protease and phosphatase inhibitors. Samples were sonicated twice for 15 seconds each in an ice-cold bath. Subsequently, the samples were incubated at 28°C in a ThermoMixer (Eppendorf) with 650 rpm agitation for 2 hours to ensure complete reduction of disulfide bonds and alkylation of cysteine thiol groups. The samples were then centrifuged at 5000x g for 10 minutes to remove cellular debris, and the supernatant was dried using a vacuum centrifuge (speedvac, Thermo). The dried samples were stored in an ultrafreezer at −80°C until further processing. The samples were resuspended in TEAB (pH = 8) and quantified using Nanodrop (Thermo Scientific). Following quantification, 300 μg of proteins were digested with 5% (w/w) trypsin at 37°C for 18 hours. Tryptic peptides were quantified using the Pierce quantitative fluorometric assay (Thermo Scientific, #23290) as per the manufacturer’s specifications. A small aliquot (1 μL) was collected to check the digestion efficiency using Orbitrap Q-exactive Plus.

### Liquid Chromatography (Evosep One)

The samples were analyzed using an HPLC instrument (Evosep One) following the manufacturer’s instructions. Tryptic peptides were loaded onto an Evosep One chromatograph (30 SPD protocol) coupled online to a timsTOF Pro mass spectrometer (Bruker). The samples were eluted from the Evotips under low pressure to the storage loop with a gradient displacement to reduce the percentage of organic buffer. Separation was achieved using a customized 40-minute gradient (30 samples/day method) at a flow rate of 2 μL/min on a reverse-phase column (4 cm x 150 μm internal diameter) packed with 3 μm C18-coated porous silica beads (PepSep, Odense, Denmark).

### Mass Spectrometry (DIA)

Label-free samples were analyzed on a timsTOF Pro mass spectrometer (Bruker) operating in PASEF mode [15] with Data Independent Acquisition (DIA). The TIMS tunnel comprises two regions: one for ion storage and another for ion mobility analysis. Trapped ion separation was performed by varying the entrance voltage in the second region. MS spectra were recorded between 100 and 1700 m/z. Suitable precursor ions for PASEF-MS/MS were dynamically selected in real-time using an advanced PASEF scheduling algorithm based on TIMS-MS survey scans. A polygonal filter was applied in the m/z and ion mobility space to identify features likely representing peptide precursors rather than singly charged ions. The quadrupole isolation window was set to 2 Th for values below 700 m/z and 3 Th for values above 700 m/z. Collision energy was incrementally adjusted depending on the increase in ion mobility: 52 eV for the first 0-19% of the ramp time, 47 eV for 19-38%, 42 eV for 38-57%, 37 eV for 57-76%, and 32 eV for the remaining part of the analysis.

### Data Processing (DIA MS)

The raw data obtained from the timsTOF PASEF were processed using the DIA-NN software, version 1.8.1. The spectral library was generated from the FASTA file of the *Mus musculus* taxonomy (Swissprot). Default parameters were used for protein data deconvolution, identifying proteins with at least two different peptides, one of which had to be unique, with a false discovery rate (FDR) < 1%. The deconvolution process in DIA-NN was executed using a script based on the instructions provided at: DIA-NN GitHub.

### Statistical Analysis (DIA MS)

Quantification values of identified proteins were log2-transformed and normalized to the median value of each column. Missing values were imputed using the “missForest” machine learning package and the statistical analysis algorithms implemented in R. Statistical calculations were performed using the unpaired t-test. For proteins to be considered differentially regulated, they had to show a p-value < 0.05 and a log2 fold change of KO vs. WT ≥ 0.263 (upregulated) or ≤ −0.263 (downregulated).

### Isobaric Labeling with TMT-16plex

Previously digested and quantified samples were dried using a speedvac and resuspended in TEAB buffer (pH = 8). The pH of all samples was checked and adjusted to pH = 8.5. Samples were labeled using the TMT-16plex kit (Lot. XI346567) according to the manufacturer’s instructions. Briefly, 20 µL of the respective TMT-16plex reagent was added to each sample and incubated at room temperature for 1 hour. A small aliquot (1 µL) of each labeled sample was tested for labeling efficiency by mixing without detergent and analyzed on an Orbitrap Q-Exactive Plus. The sample quantities in each tube were adjusted to ensure equal proportions of the 16 experimental groups. After labeling, each sample was acidified with 2% (v/v) formic acid, vigorously agitated, and centrifuged for 10 minutes at 14000x g. The supernatant was collected and dried using a vacuum centrifuge.

### Enrichment of Phosphopeptides and N-glycopeptides

Phosphopeptides were enriched using the TiO2 method, as previously reported [16]. TMT-16plex labeled samples were resuspended in a loading solution containing 80% acetonitrile (ACN), 5% trifluoroacetic acid (TFA), and 1 M glycolic acid. Subsequently, samples were incubated for 30 minutes with titanium dioxide (TiO2) particles at a ratio of 6:1 (sample (μg): TiO2 (μg)) under vigorous agitation. To maximize phospho-enrichment efficiency, the loading solution containing non-resin-bound phosphopeptides from the previous step was transferred to a second tube and incubated again with TiO2 particles at a ratio of 3:1 (sample (μg): TiO2 (μg)) for 30 minutes.

Particles were washed with wash buffer 1 containing 80% ACN and 1% TFA (v/v) for 10 seconds and centrifuged to separate TiO2 particles. The supernatant was collected and transferred to a new tube containing unmodified peptides. Particles were washed again with wash buffer 2 containing 10% ACN and 0.2% TFA (v/v) for 10 seconds and centrifuged again to separate TiO2 particles. The supernatant was collected and transferred to a new tube containing peptides from the second wash. TiO2 particles were dried for 5 minutes in a vacuum centrifuge (Eppendorf, Germany). Modified peptides retained on TiO2 particles were eluted with 100 μL elution buffer containing 25% ammonia (v/v) and pH = 11.3 under vigorous agitation for 15 minutes. The solution was then centrifuged for 1 minute at 14000xg, and the supernatant was filtered with a C8 resin-containing pipette tip. This process was repeated once more using 30 μL elution buffer with TiO2 particles. Modified peptides were eluted from the C8 resin with 5 μL of a solution containing 30% ACN (v/v) and transferred to the eluted peptide tube. After this step, the solution containing eluted peptides was dried using a vacuum centrifuge (Eppendorf, Germany). Unmodified peptides were purified using commercial Waters™ Oasis HLB Cartridges columns, and after desalting, samples were dried using a vacuum centrifuge.

### Deglycosylation Reaction

For deglycosylation, modified peptides were resuspended in 150 μL of a solution containing 20 mM TEAB at pH 7.5. 1 µL of PNGase F enzyme and 0.5 µL of Sialidase A were added to the samples. Samples were incubated overnight at 37°C. After incubation, 5 µL of 100% formic acid (v/v) and 5 µL of 10% TFA (v/v) were added to the sample to stop the reaction. The pH of the samples after this process was checked and found to be pH < 2.

### Pre-fractionation of Samples by Basic pH Reverse-Phase Liquid Chromatography

TMT16-plex labeled proteome samples were pre-fractionated using an ultra-performance liquid chromatography system. Peptides were separated into 60 fractions and combined into 20 final fractions sequentially. Mobile phases used were solution A (ammonium formate pH 9.3) and B (80% ACN (v/v) and 20% solution A). The chromatographic method consisted of a flow rate of 5 µL per minute and the following elution gradient: (i) 0-14 min 2% ACN; (ii) 14-20 min 2-10% ACN; (iii) 20-85 min 10-40% ACN; (iv) 85-117 min 40-50% ACN; (v) 117-122 min 50-95% ACN; (vi) 122-132 min 95% ACN; (vii) 132-133 min 2% ACN. After fractionation, samples were dried using a vacuum centrifuge.

Samples containing modified peptides were separated into 36 fractions and combined into 12 final fractions sequentially. The chromatographic method consisted of a flow rate of 5 µL per minute and the following elution gradient: (i) 0-14 min 2% ACN; (ii) 14-17 min 2-12% ACN; (iii) 17-49 min 12-40% ACN; (iv) 49-59 min 40-95% ACN; (v) 59-69 min 95% ACN; (vi) 70-86 min 2% ACN. After fractionation, samples were dried using a vacuum centrifuge.

### Mass Spectrometry Analysis (DDA)

Samples of unmodified (proteome) and modified peptides (phosphoproteome and N-glycoproteome) were resuspended in 0.1% formic acid (v/v) solvent A and analyzed using an UPLC with nanoflow EASY-nLC (Thermo), coupled to an Exploris 480 mass spectrometer (Thermo). The pre-column was approximately 3 cm long, with an internal diameter of 100 μm, packed with C18 AQ Reprosil-Pur resin with 5 μm diameter particles. The analytical column (18 cm x 75 μm) was packed with C-18 resin with 3 μm diameter particles. The chromatographic gradient used was: (i) 8-25% solvent B (95% ACN (v/v) and 0.1% formic acid (v/v)) over 69 minutes; (ii) 25-45% solvent B in 15 min; (iii) 45-95% solvent B; (iv) 95% solvent B for 5 minutes; (v) 95-1% solvent B for 1 minute; (vi) 95-1% solvent B for 5 minutes at a flow rate of 300 nL/min. The mass spectrometer (Orbitrap Exploris 480) was operated in positive polarity and data-dependent acquisition (DDA) mode. Peptides were analyzed in MS1 with a range of 350-1600 m/z. Parent ions were accumulated up to 3×10^6^ or for 100 ms, whichever occurred first (MS1 AGC target settings). Parent ions were resolved with a resolution of 120000 FWHM (full width at half maximum). The top 20 most intense ions (Top 20) were selected in the quadrupole with an isolation window of 0.7 Th for higher energy collision dissociation (HCD), with a normalized collision energy of 33%. Injection time for fragment ions was 20 ms or accumulation of 10^6^ ions, whichever occurred first (MS2 AGC target settings). Fragment ions were resolved with 17,500 FWHM at 200 m/z and their precursor ions were included in the dynamic exclusion list per second.

Samples containing modified peptides (phosphoproteome and N-glycoproteome) were simultaneously analyzed using the Orbitrap Fusion Lumos. The chromatographic gradient used was: (i) 2-8% solvent B (95% ACN (v/v) and 0.1% formic acid (v/v)) for 5 minutes; (ii) 8-28% solvent B for 85 minutes; (iii) 28-40% solvent B for 15 minutes; (iv) 40-95% solvent B for 3 minutes; (v) 95% solvent B for 5 minutes; (vi) 95-0% solvent B for 3 minutes at a flow rate of 300 nL/min. The mass spectrometer (Orbitrap Fusion Lumos) was operated in positive polarity and DDA mode. Peptides were analyzed in MS1 with a range of 350-1600 m/z. Parent ions were accumulated up to 3×10^6^ or for 100 ms, whichever occurred first (MS1 AGC target settings). Parent ions were resolved with a resolution of 120000 FWHM. The top 20 most intense ions (Top 20) were selected in the quadrupole with an isolation window of 0.7 Th for higher energy collision dissociation (HCD), with a normalized collision energy of 34%. Injection time was 50 ms or accumulation of 10^6^ ions, whichever occurred first. Fragment ions were resolved with 17,500 FWHM at 200 m/z and their precursor ions were included in the dynamic exclusion list per second. Raw spectra (.raw) were viewed with Xcalibur v3.0 (Thermo).

### Bioinformatics Analysis (DDA MS)

The raw data (.raw) was used for protein database searches using MaxQuant 2.1.0.0 against a FASTA file downloaded in December 2020 from UniProtKB/SwissProt *Mus musculus* (17,492 entries). A list of common contaminants was included in the search parameters, with a tolerance of 20 ppm for the first search and 4.5 ppm for the main search. MS2 quantification was chosen, and TMT-16plex labeling was configured for isobaric labeling quantification. Trypsin was selected as the enzyme, allowing for a maximum of two missed cleavage sites. Carbamidomethylation (Cys) was chosen as a fixed modification, while oxidation (Met) and acetylation (N-terminal) were set as variable modifications. For modified peptide samples, phosphorylation (STY) and deamidation (N) modifications were enabled. “Match between runs” was activated with a time window of 0.7 min and alignment within 20 min. For peptide identification, a minimum of 1 unique peptide was required. The false discovery rate (FDR) was set to <1%, and a reverse decoy database was used to determine FDR. Perseus was used to transform protein abundance values to log2 scale and normalize abundance values (log2) for each experimental group. Statistical analyses were performed using t-test (p < 0.05) in the statistical program “R”.

### Measurement of exocytosis rate using FM1-43 probe

Synaptic vesicle staining was performed using the fluorescent probe FM1-43 [17, 18]. Studies were performed according to the previously described by [19]. Brain slices containing the striatum and substantia nigra were incubated in high-potassium artificial cerebrospinal fluid (aCSF) containing (120mM NaCl, 60 mM KCl, 1.3 mM MgCl2, 2.5 mM CaCl2, 1.25 mM NaH2PO4, 26 mM NaHCO3, and 11 mM glucose, gassed with 95% O2/5% CO2 and 8 μM FM1-43) for 1.5 minutes to promote dye uptake. Slices were then washed in normal aCSF containing (120mM NaCl, 3.5 mM KCl, 1.3 mM MgCl2, 2.5 mM CaCl2, 1.25 mM NaH2PO4, 26 mM NaHCO3, and 11 mM glucose, gassed with 95% O2/5% CO2) for 2 minutes, followed by perfusion with 1 mM ADVASEP-7 for 1 minute to remove excess extracellular dye. This was followed by another wash in normal saline for 2 minutes, a second perfusion with 1 mM ADVASEP-7 for 1 minute, and a final wash in normal saline for 20 minutes to ensure complete removal of unbound dye. Fluorescence imaging was performed using a Leica DM2500 fluorescence microscope equipped with a 63× water immersion objective (0.94 NA) and coupled to a 12-bit CCD Micromax camera (Leica DFC345FX). Image acquisition and visualization were conducted using the Leica Application Suite 4.0 software. Excitation light was provided by an HXP R120/45C-VIS lamp and passed through a 505/530 nm filter set to isolate the specific fluorescence spectrum (λex= 488 nm; λem = 505–520 nm). All imaging parameters were kept constant across control and experimental conditions. Fluorescence analysis was performed using ImageJ and CellProfiler software. Following image alignment, mean fluorescence intensity was quantified for each time point. To account for variability in baseline fluorescence among samples, data were normalized by expressing fluorescence values as a percentage of the initial intensity (minute 0). This normalization enabled reliable comparisons of fluorescence decay across samples, minimizing variation due to differences in initial signal levels.

### Statistical analysis of exocytosis rate measurement

To assess differences in the dynamics of decay curves between experimental conditions, linear regression models were fitted for each group separately. An analysis of covariance (ANCOVA) was then performed using linear models including the interaction between time and group (model: lm(Decay ∼ Time * Group)) to evaluate whether the slope of the regression line differed significantly between groups. Pairwise comparisons were conducted by fitting models restricted to two groups at a time. Statistical significance was assessed based on the interaction term (Time:Group), with *p*-values obtained from model summaries. Values of *p* < 0.05 were considered statistically significant.

### High-Performance Liquid Chromatography analysis

To quantify the neurotransmitters of interest, we employed a method similar to that described previously [20]. The brain regions were resuspended in a deproteinization solution containing 0.2 M perchloric acid and 3 mM cysteine. The samples were then sonicated in an ice bath and centrifuged at 15,000 x g for 15 minutes at 4°C. The supernatant was collected and analyzed using a high-performance liquid chromatography system equipped with a fluorescence detector (HPLC-FD, Shimadzu). A C18 chromatography column (250 x 4.60 mm, 5μ; Phenomenex) was used as the stationary phase. The mobile phase consisted of a solution containing 12 mM CH3COONa and 0.26 mM EDTA.Na2 (pH = 3.5). The instrument was run under isocratic conditions with a flow rate of 0.5 mL/min for 30 minutes. Neurotransmitter detection was performed using excitation and emission wavelengths of λex = 279 nm and λem = 320 nm, respectively, with high sensitivity and a gain of 4x.

### Statistical analysis of neurotransmitters quantifications

For comparisons between two independent experimental groups, when data followed a normal distribution and variances were comparable, a two-tailed unpaired Student’s *T*-test was applied. A *p*-value < 0.05 was considered statistically significant. Data are presented as mean ± standard error of the mean (SEM).

### Behavioral tests Open field Test

The Open Field Test was employed to investigate the locomotor activity of the mice [21]. The experiment was conducted in a white box (50 cm × 50 cm × 30 cm) (Insight®, São Paulo, Brazil) and involved placing each mouse in the center of the box, allowing them to explore the space freely for a period of 30 minutes. The total distance traveled during this exploration was meticulously recorded using the Ethovision software developed by Noldus Information Technology. The apparatus was cleaned between test sessions with a 10% ethanol solution.

### Elevated plus maze test

The elevated plus maze consists of two open arms and two closed arms. During the test, each mouse was placed in the center of the platform and allowed to freely explore the maze for 5 minutes. The number of entries into the open and closed arms, as well as the percentage of time spent in the open arms, were measured [22]. The apparatus was cleaned between test sessions with a 10% ethanol solution.

### Marble burying test

Mice were individually placed in a polycarbonate box (30 x 40 x 12 cm) with sawdust for five minutes to habituate. They were then removed and placed in their regular housing for 30 minutes. After this period, the mice were reintroduced to the same box containing 20 glass beads (1.5 cm in diameter) distributed equidistantly for 15 minutes. After this period, the animals were removed from the box, and the number of buried beads was counted. Beads were considered buried if at least two-thirds of their area were covered by sawdust.

### Y maze

The Y Maze Test evaluates the short-term memory capacity of rodents. This test checks if the rodent retains the memory of the space it has just explored. The experiments were conducted in a sound-attenuated environment with low-intensity light. The mice was placed at the end of one arm of the Y maze (central position of the apparatus) and was free to explore the three arms for 8 minutes, with the first 2 minutes for habituation and the remaining 6 minutes for testing. The number of alternations between the three arms (A, B, and C) was counted during the test period [23]. The apparatus was cleaned between test sessions with a 10% ethanol solution.

### Novel Object Recognition test (NOR)

The Novel Object Recognition Test was performed as previously described [24]. The test consisted of three phases: habituation, familiarization (also called training), and testing. During the habituation phase, the animals were exposed for 5 minutes to an opaque acrylic arena (approximately 38 cm wide, 38 cm long, and 15 cm high). Twenty-four hours after habituation, the animals were placed back in the center of the same environment and monitored for 10 minutes in the presence of two identical copies of the same object (objects A1 and A2, positioned on opposite sides, away from the walls). On the third day (training), the animals were placed back in the center of the same environment and monitored for 10 minutes in the presence of objects A1 and A2. On the fourth day, a new object was introduced to the animals, and the exploration time of the new object (A3) and the familiar object (A1) was recorded for 5 minutes using a camera system and manually analyzed with stopwatches. The objects used were two small plastic bottles of the same color, size, and texture, and a Lego toy similar in height and diameter to the bottle. Objects A1 and A2 were 17 cm tall and 3-5 cm in diameter (varying along the object), and the novel object B was 13.5 cm tall and 5 cm in diameter. After each test session, the location and objects were cleaned with 10% ethanol. Additionally, whenever placed in the arena, the animals were oriented opposite to the objects. All sessions were conducted under 303 lux of light intensity.

### Behavioral statistical analysis

All behavioral results were analyzed by a second person and the statistical analyses were performed using GraphPad Prism 8. The choice of statistical test was based on the experimental design and data structure. For behavioral tests involving repeated measures over time, such as the Open Field test, a mixed-effects model (REML method) was applied. Time (3, 6, and 12 months) was treated as a within-subjects fixed effect, and experimental group (WT and KO) as a between-subjects fixed effect. When appropriate, Geisser–Greenhouse correction was applied to account for violations of sphericity. Interactions between time and group were also assessed. For tests performed at a single time point, such as the Y maze and NOR unpaired two-tailed student’s *t*-tests were used for normally distributed data. Data are presented as mean ± standard error of the mean (SEM). Differences were considered statistically significant when *p* < 0.05.

## RESULTS

### PTMomics Analysis of the Nigrostriatal Pathway in Mrgprd Deficient Mice

To understand the molecular alterations resulting from the absence of the MrgD receptor in the nigrostriatal pathway, proteome, phosphoproteome, and N-glycoproteome analyses were performed on the substantia nigra (SN) and striatum (ST) of WT and KO mice (Figure 1A). In total, 5,407 proteins were quantified in the nigrostriatal pathway using DIA-PASEF, of which 148 proteins were positively regulated and 154 negatively regulated in the SN, while 56 proteins were positively regulated and 110 negatively regulated in the ST (Figure 1B). Proteins, phosphosites, and glycosites with p < 0.05 and abundance ± 0.263 in log2 were considered regulated. Additionally, we identified 19,209 phosphosites (STY) and 5,301 N-glycosites in the nigrostriatal pathway. Of these, 143 phosphosites were positively regulated and 43 negatively regulated in the SN, while 5 phosphosites were positively regulated and 7 negatively regulated in the ST. In the N-glycoproteome, we observed 47 positively regulated glycosites and 14 negatively regulated glycosites in the SN, and 1 positively regulated glycosite and 5 negatively regulated glycosites in the ST. For analysis, only phosphosites and N-glycosites with a phosphorylation/glycosylation probability ≥75% (Class 1) were considered (Figure 1C).

**Figure 1.**
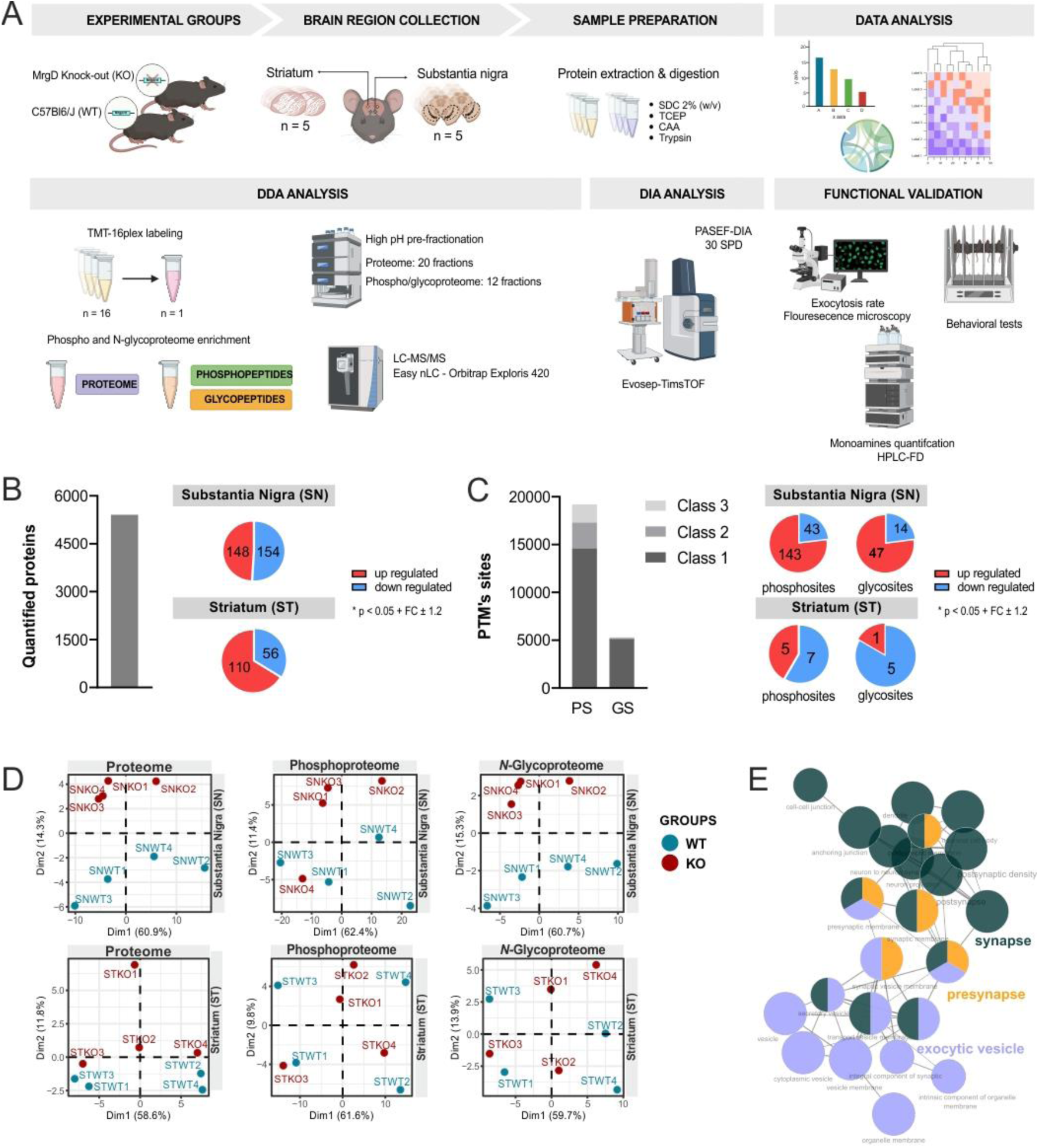
Proteomics analysis of the nigrostriatal pathway of Mrgprd deficient mice. A) Proteomics workflow of the PTMomics analysis. B) Number of quantified proteins and the number of regulated proteins in the substantia nigra (SN) and striatum (ST). C) Number of phospho- and glycol-sites and regulated sites in the SN and ST. D) Principal component analysis of the proteome, phosphoproteome and N-glycoproteome datasets in the SN and ST. E) GO analysis (Celular Component) of all regulated proteins, phosphoproteins and glycoprotein in both tissues. Abbreviations: PS = Phosphosites; GS = Glycosites.

Principal component analysis (PCA) showed a clear separation between WT and KO groups in the proteome (PC1 = 60.9%, PC2 = 14.3%) and glycoproteome (PC1 = 60.7%, PC2 = 15.3%) of the substantia nigra (SN). In the SN phosphoproteome, although one of the KO samples clustered closer to the WT, a reasonable separation between the groups was still observed (PC1 = 62.4%, PC2 = 11.4%). In the striatum (ST), the proteome showed moderate separation (PC1 = 58.6%, PC2 = 11.8%), while phosphoproteome (PC1 = 61.6%, PC2 = 9.8%) and glycoproteome (PC1 = 59.7%, PC2 = 13.9%) data showed greater overlap between groups, suggesting a lesser impact of the absence of the MrgD receptor in this tissue. Overall, the results indicate that deletion of the MrgD receptor causes more pronounced molecular alterations in the substantia nigra than in the striatum (Figure 1D).

To investigate the proteins impacted by Mrgprd gene deletion in the nigrostriatal pathway, a Gene Ontology analysis based on terms associated with cellular components was performed, using regulated proteins, phosphoproteins, and glycoproteins in both tissues. Enrichment showed that regulation in the nigrostriatal pathway was predominantly related to exocytic vesicles, pre-synapse, synapse, and post-synapse along the nigrostriatal pathway (Figure 1E).

To evaluate which biological processes (BP) the negatively regulated proteins in the substantia nigra were related to, enrichment of BPs was performed using the Database for Annotation, Visualization and Integrated Discovery (DAVID). In the substantia nigra, negatively regulated proteins showed significant enrichment in BPs related to calcium ion transport, synaptic vesicle exocytosis, chemical synaptic transmission, regulation of long-term synaptic plasticity, locomotor behavior, cAMP response, protein phosphorylation, and modulation of voltage-dependent calcium channel activity (Figure 2A). Among these proteins, SNARE complex proteins stand out, such as Synaptotagmin 1 (Syt1), an essential calcium sensor that induces the fusion of synaptic vesicles with the plasma membrane [25] and Syntaxin 1 (Stx1a), another crucial component of the SNARE complex, fundamental for controlled neurotransmitter release [26]. Complexin 2 (Cplx2), which stabilizes the SNARE complex and regulates vesicular fusion in a calcium-dependent manner [27], and Dynamin 2 (Dnm2), a GTPase that plays a crucial role in membrane cleavage during vesicle endocytosis, being necessary for the recycling of these synaptic vesicles [28], were also negatively regulated (Figure 2B). Stx1a and Cplx2 proteins showed negative regulation in both mass spectrometry techniques used (Table S4 and Table S5). The negative regulation of these proteins indicates that the absence of the MrgD receptor appears to alter the dynamics of synaptic release in the substantia nigra, compromising the efficiency of neurotransmission in this region.

**Figure 2.**
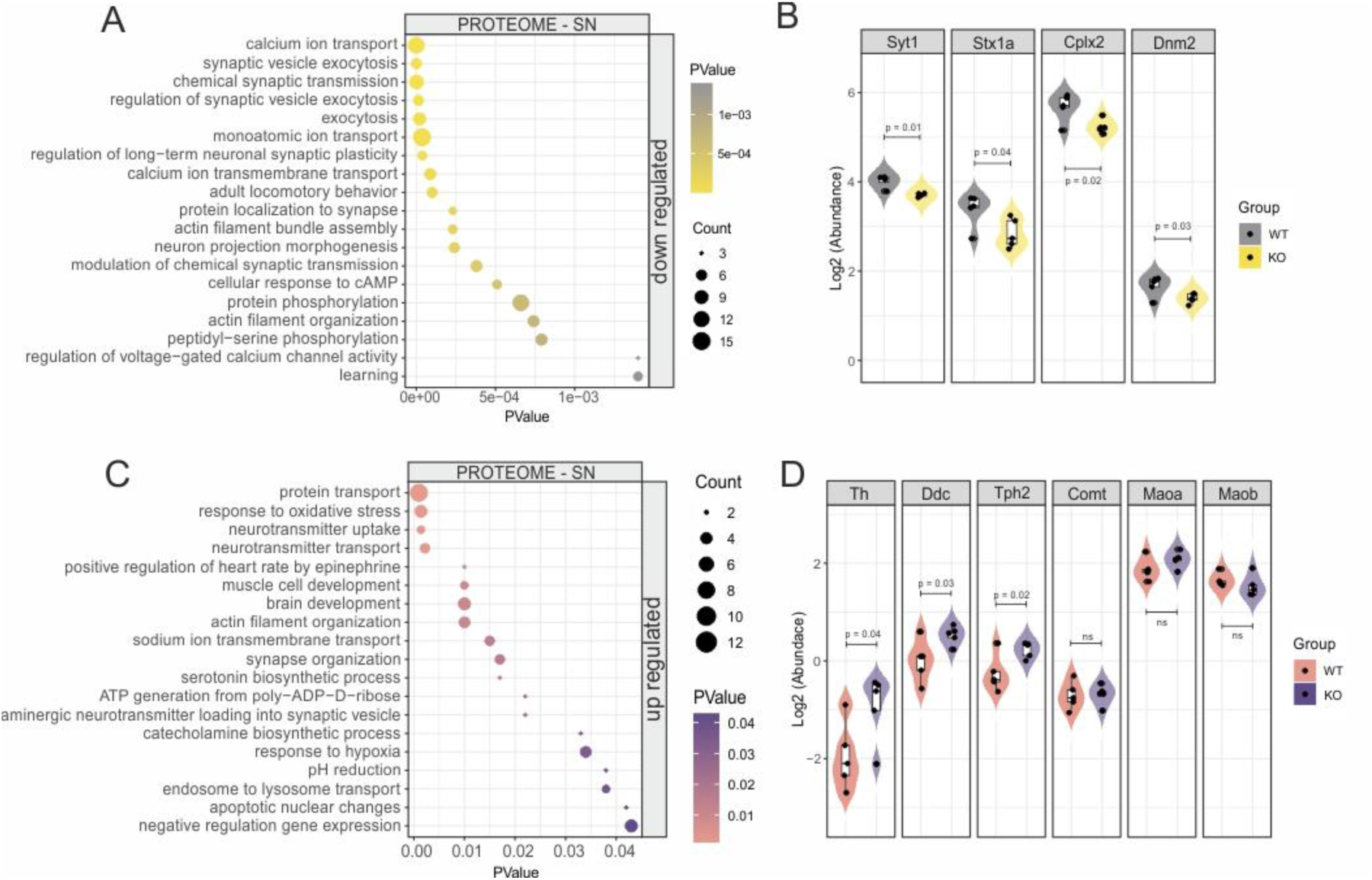
Functional enrichement in the substantia nigra of Mrgprd deficient mice. A) Biological process (BP) enrichment of the downregulated proteins in the SN. B) SNARE proteins being downregulated in the SN. C) BP enrichment of the upregulated proteins in the SN. D) Enzymes associated with monoamine metabolism in the SN.

Functional enrichment using the positively regulated proteins showed that MrgD receptor deficiency led to the positive regulation of proteins involved in oxidative stress response, hypoxia, neurotransmitter uptake and transport, epinephrine-mediated heart rate regulation, and the catecholamine biosynthetic process, as well as the loading of aminergic neurotransmitters into synaptic vesicles in the substantia nigra (Figure 2C). The positive regulation of these pathways suggests that the absence of the MrgD receptor appears to trigger adaptive responses in the substantia nigra, potentially as a compensatory mechanism against synaptic dysfunction. The increase in the expression of proteins associated with neurotransmitter biosynthesis and transport may indicate an attempt to maintain neurochemical and functional homeostasis.

Notably, essential enzymes for catecholamine synthesis in the substantia nigra, such as Tyrosine Hydroxylase (Th), responsible for the conversion of tyrosine to L-DOPA, the rate-limiting step of catecholamine biosynthesis [29]; DOPA Decarboxylase (Ddc), which catalyzes the conversion of L-DOPA to dopamine [30]; and Tryptophan Hydroxylase 2 (Tph2), involved in the hydroxylation of tryptophan for serotonin formation, which also plays modulatory roles in dopaminergic circuits [31] were positively regulated. In contrast, no significant changes were observed in the abundance of enzymes associated with catecholamine metabolism, such as Catechol-O-Methyltransferase (Comt), which mediates catecholamine methylation, and Monoamine Oxidase A (Maoa) and Monoamine Oxidase B (Maob) enzymes, responsible for the oxidative degradation of neurotransmitters such as dopamine, norepinephrine, and serotonin [32] (Figure 2D).

The Ddc protein was negatively regulated in both mass spectrometry techniques used (Table S4 and Table S5). Together, the proteome-level data indicate a decrease in synaptic vesicles accompanied by an increase in monoamine production in the substantia nigra.

To investigate possible diseases associated with MrgD receptor deletion in the substantia nigra, a disease enrichment analysis was performed based on the Comparative Toxicogenomics Database (CTD) with the regulated proteins in the substantia nigra. The regulated proteins related to neurotransmission and monoamine production are intimately related to Parkinson’s disease, neurodegeneration, Schizophrenia, bipolar disorder, and attention deficit hyperactivity disorder (Figure S1). Therefore, the MrgD receptor appears to play a relevant role in maintaining the functional integrity of the substantia nigra, and its absence seems to contribute to molecular alterations associated with neuropsychiatric and neurodegenerative disorders.

Considering the central role of the post-translational modifications in regulating stability, subcellular localization, protein-protein interaction, and function of proteins involved in neurotransmission, we performed the N-glycoproteome and phosphoproteome analysis in the substantia nigra of Mrgprd knockout mice. The differentially regulated phosphoproteins and glycoproteins were mainly associated with processes such as learning, modulation of chemical synaptic transmission, calcium ion transport, synaptic vesicle endocytosis, memory, neurotransmitter secretion, locomotor behavior, among others (Figure 3A). The enrichment of kinases potentially responsible for these modifications indicated the involvement of cyclic AMP and calcium-dependent protein kinases (PKA and PKC) (Figure 3B). These findings suggest that the absence of the MrgD receptor results in altered activity of basophilic kinases, such as PKA and PKC in the substantia nigra, which may indicate the participation of MrgD in the activation of intracellular signaling pathways mediated by Gs or Gq type G proteins, which are important for neurotransmission, neuronal plasticity and modulation of motor behavior in SN [33].

**Fig 3.**
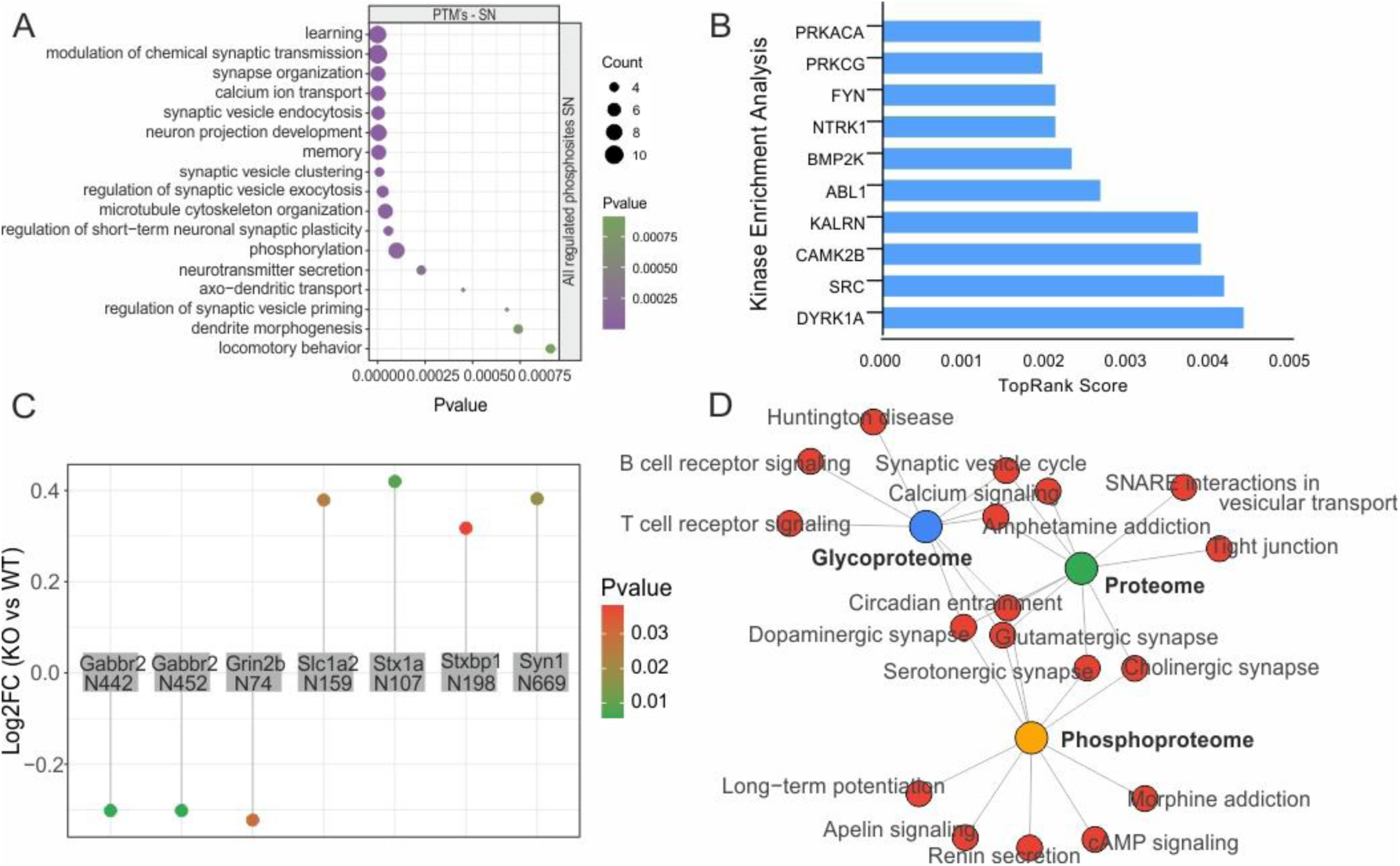
Integration of the multiomics data. A) BP enrichment using the phosphosites and glycosites regulated in the SN. B) Kinase enrichment analysis using the regulated phosphoproteins in SN. C) Regulated glycosites in the SN. D) Simultaneos KEGG pathways enriched in each proteome level in the SN. The graph was performed using “igraph” in R.

Among the differentially N-glycosylated proteins identified, several with essential functions in neurotransmission stand out. Among them, a reduction in the glycosylation of two sites of the GABAB receptor subunit (Gabbr2) at Asn-442 and Asn-452 residues was observed, which may reflect alterations in inhibitory GABAergic signaling. The NMDA receptor subunit (Grin2b), crucial for excitatory glutamatergic transmission and synaptic plasticity, also showed decreased glycosylation at the Asn-74 residue. On the other hand, the glutamate transporter (Slc1a2) showed increased glycosylation at the Asn-159 residue. Similarly, SNARE complex proteins Stx1a, syntaxin-binding protein (Stxbp1), synapsin 1 (Syn1) showed an increase in glycosylation at Asn-107, Asn-198, and Asn-669 residues, respectively (Figure 3C). These proteins are directly involved in glutamate uptake, synaptic vesicle fusion, and neurotransmitter release, suggesting a possible modulation of synaptic activity via glycosylation in the substantia nigra without the presence of the MrgD receptor.

The integration of data from proteomic, phosphoproteomic, and N-glycoproteomic analyses provided a comprehensive view of the signaling pathways modulated in the substantia nigra. It was observed that the dopaminergic synapse, glutamatergic synapse, and circadian cycle pathways showed simultaneous regulation at all three molecular levels. Additionally, alterations were identified in calcium signaling pathways and the synaptic vesicle cycle at the proteome and N-glycoproteome levels, while pathways related to serotonergic and cholinergic synapses showed regulation at both proteome and phosphoproteome levels. Specific alterations were also observed in immunological pathways, such as B and T cell receptor signaling in the glycoproteome; in long-term potentiation and cAMP signaling in the phosphoproteome; and in SNARE interaction in vesicular transport in the proteome (Figure 3C). Together, the data indicate the modulation of multiple synaptic and intracellular signaling pathways in the substantia nigra, which can affect different neurotransmission pathways.

### Exocytosis Rate Measurement in the Nigrostriatal Pathway in Mrgprd Deficient Mice

To confirm whether synaptic transmission was functionally compromised in KO mice, the rate of synaptic vesicle exocytosis was measured in the substantia nigra and striatum regions using the fluorescent probe FM1-43. In agreement with proteomic data, a smaller fluorescence decay was observed in the tissues of KO animals compared to controls in both the substantia nigra (Figure 4A) and the striatum (Figure 4B), indicating a reduction in the rate of synaptic exocytosis in these regions. Moreover, a significant increase in norepinephrine (NE), epinephrine (E), dopamine (DA), and serotonin (5-HT) levels was observed in KO mice in the substantia nigra (Figure 4C). On the other hand, in the striatum, there was a reduction in dopamine levels, with no changes in NE and 5-HT levels in KO mice compared to WT (Figure 4D). These data, combined with the negative regulation in abundance and post-translational modifications observed in SNARE complex components, indicate an impairment in dopaminergic neurotransmission between the two regions. These results reinforce the hypothesis that the absence of the MrgD receptor directly affects the mechanisms of neurotransmission in the nigrostriatal pathway.

**Figure 4.**
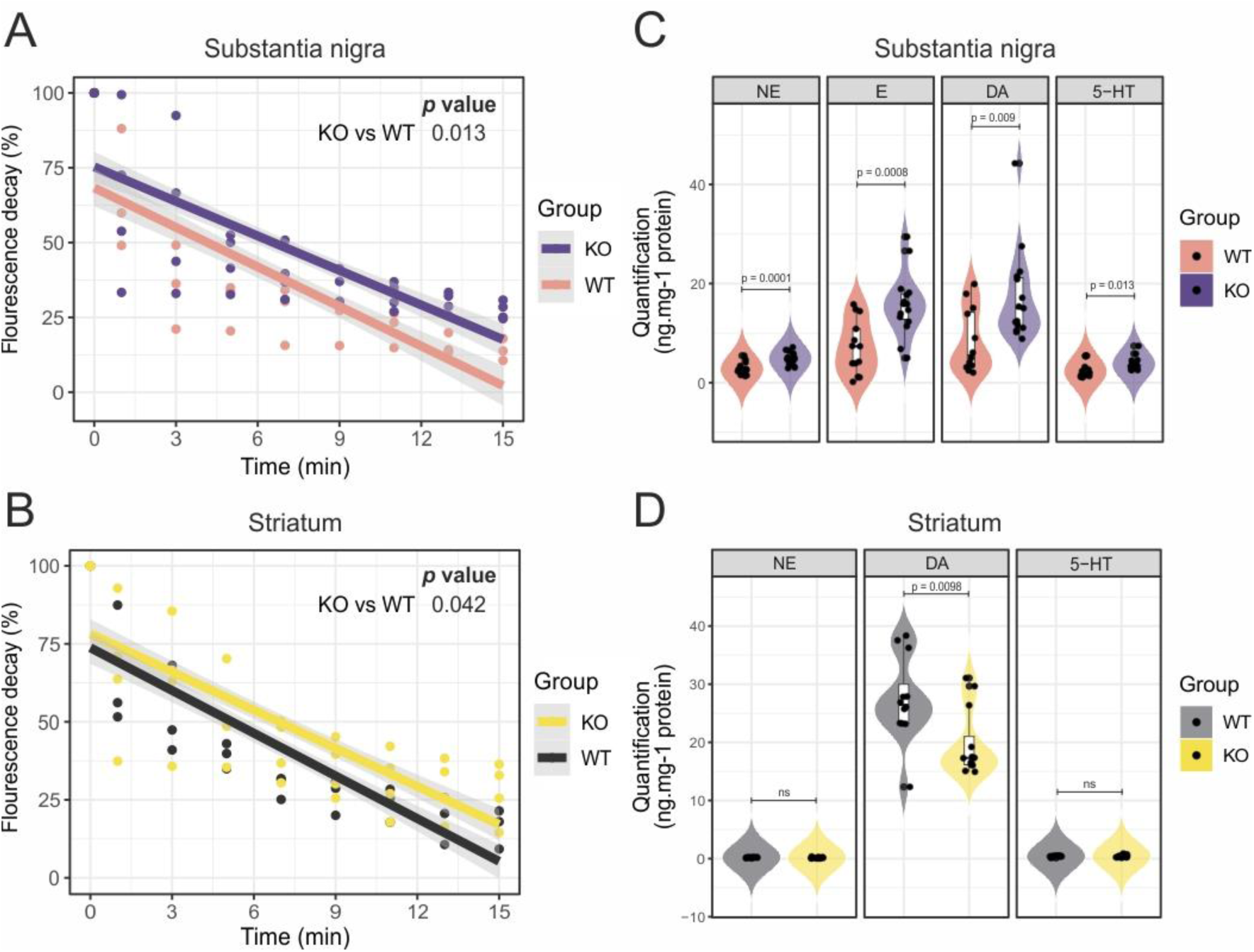
Exocytosis rate measurement in the nigrostriatal pathway. A) Measurement of exocytosis rate using the fluorescent probe FM1-43 in SN and B) in the ST. C) Quantification of monoamines using HPLC coupled to fluorescent detector in the SN and (D in the ST.

### Pharmacodynamic Specificity of MrgD in Synaptic Transmission

To evaluate the role of the MrgD receptor in synaptic vesicle exocytosis, control mice (WT) were treated with the best-characterized receptor agonists, β-alanine [10^-4^ M] (WT + β-alanine) and alamandine [10^-7^ M] (WT + Ala), for 15 minutes in tissue slices from the substantia nigra and striatum and performed the measurement of the exocytosis rate in these tissues. Interestingly, the stimulation with β-alanine did not produce significant changes in synaptic exocytosis, either in the substantia nigra (Figure 5A) or in the striatum (Figure 5B). In contrast, treatment with alamandine resulted in a significant increase in the exocytosis rate in both regions (Figure 5A and 5B).

**Figure 5.**
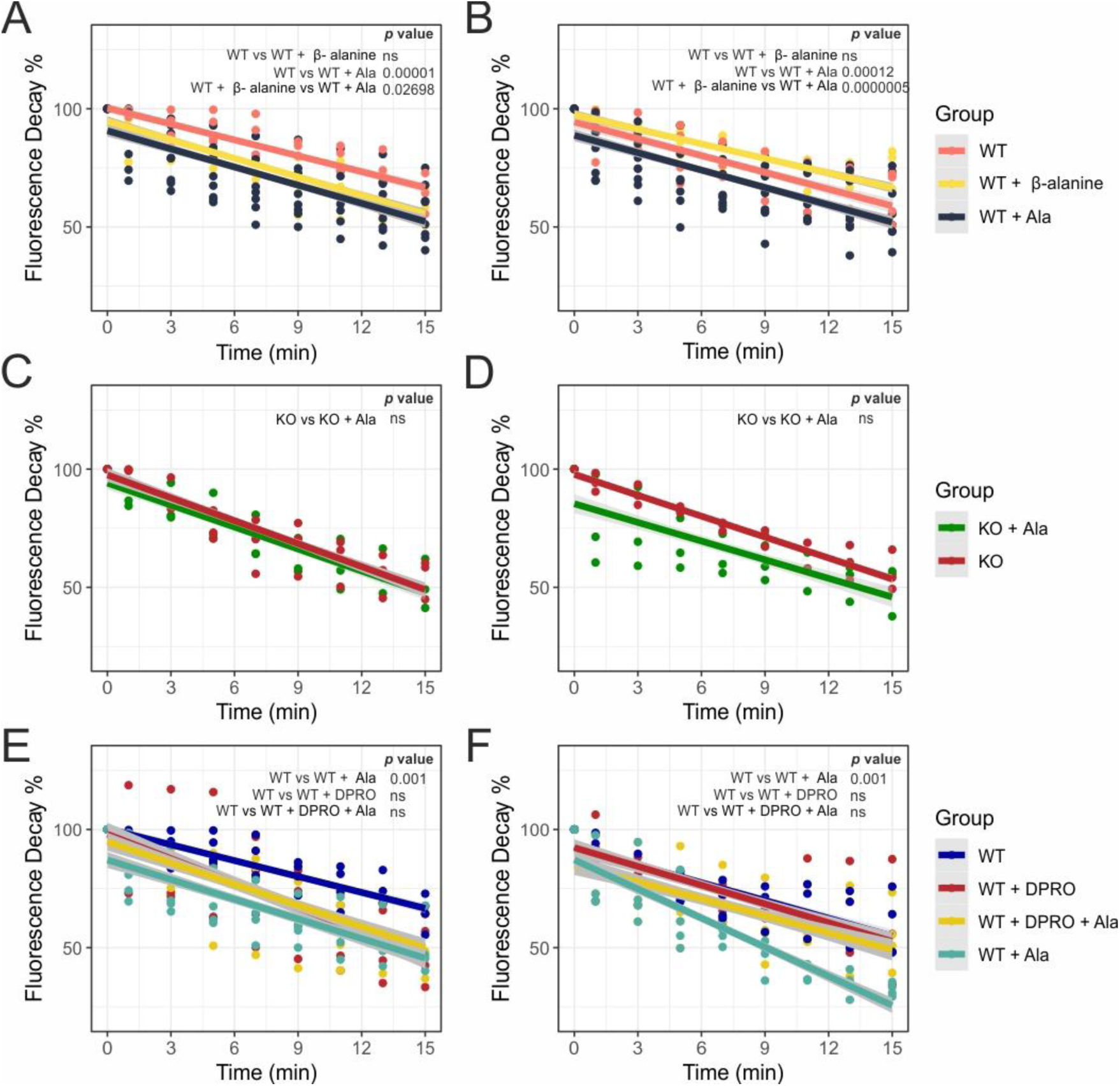
Pharmacodynamic Specificity of MrgD in Synaptic Transmission. Measurement of exocytosis rate in the A) SN of wild type (WT), WT stimulated with β-alanine (WT + β-alanine) and WT + Alamandine (WT + Ala) and in the B) ST. C-D) measurement of exocytosis rate in the SN and ST using Mrgprd-knockout mice (KO) and KO stimulated with Alamandine (KO + Ala). E-F) Measurement of exocityosys rate in the SN and ST using WT mice, WT + Ala, WT stimulated with MrgD partial antagonist DPRO (WT + DPRO) and WT + Ala + DPRO.

To confirm whether vesicle release was indeed being modulated by the MrgD receptor, KO mice were subjected to alamandine [10^-7^ M] treatments for 15 min (KO + Ala) in tissue slices from the substantia nigra and striatum. Statistical analysis comparing the models between group and time indicated that there was no significant difference in the slope of the curves between the alamandine-treated KO and control KO groups, either in the substantia nigra or in the striatum (Figure 5C and 5D).

To more comprehensively evaluate the pharmacological modulation of the MrgD receptor on synaptic exocytosis, control (WT) mice were distributed into four experimental groups: i) WT without treatment; ii) WT treated with the partial antagonist DPRO [10^-6^ M] for 15 min (WT + DPRO); iii) WT pretreated with DPRO for 15 min [10^-6^ M] and subsequently stimulated with alamandine [10^-7^ M] for 15 min (WT + DPRO + Ala); and iv) WT treated with alamandine alone [10^-7^ M] for 15 min (WT + Ala). As expected, no significant difference in vesicle release was observed between the WT vs WT + DPRO groups, nor between WT vs WT + DPRO + Ala; however, a significant difference was detected between the WT vs WT + Ala groups in substantia nigra and striatum (Figure 5E and 5F).

These data suggest that the MrgD receptor modulates synaptic exocytosis in an agonist-selective manner, being responsive to alamandine but not to β-alanine. This selectivity may stem from differences in ligand binding, functional efficacy, or the activation of distinct downstream signaling pathways, underscoring the complex pharmacodynamics of the MrgD receptor. Furthermore, the lack of response to alamandine in MrgD knockout animals indicates that this receptor is essential for alamandine-induced modulation of synaptic vesicle release. Consistently, treatment with the partial MrgD antagonist D-Pro was able to partially inhibit the synaptic vesicle release triggered by alamandine, further supporting the receptor’s role in mediating this effect.

### MrgD Deficiency Leads to Motor Hyperactivity and Compulsive-Like Behavior Without Memory Impairment

To further explore the functional consequences of the molecular changes observed in the nigrostriatal pathway, we conducted a series of behavioral tests. These tests were designed to evaluate anxiety-like behavior, memory, and locomotor activity, all of which are likely to be influenced by the altered neurotransmitter levels and synaptic transmission in this pathway. Surprisingly, MrgD-deficient mice traveled a greater total distance in the open field test (OFT) (Figure 6A). This increased activity persisted up to 12 months of age. The percentage of entries into the open arm and time spent in the open arm in the elevated plus maze also showed no significant differences (Figure 6B and 6C). However, MrgD-deficient mice buried significantly more marbles than wild-type mice across all ages assessed (Figure 6D).

**Figure 6.**
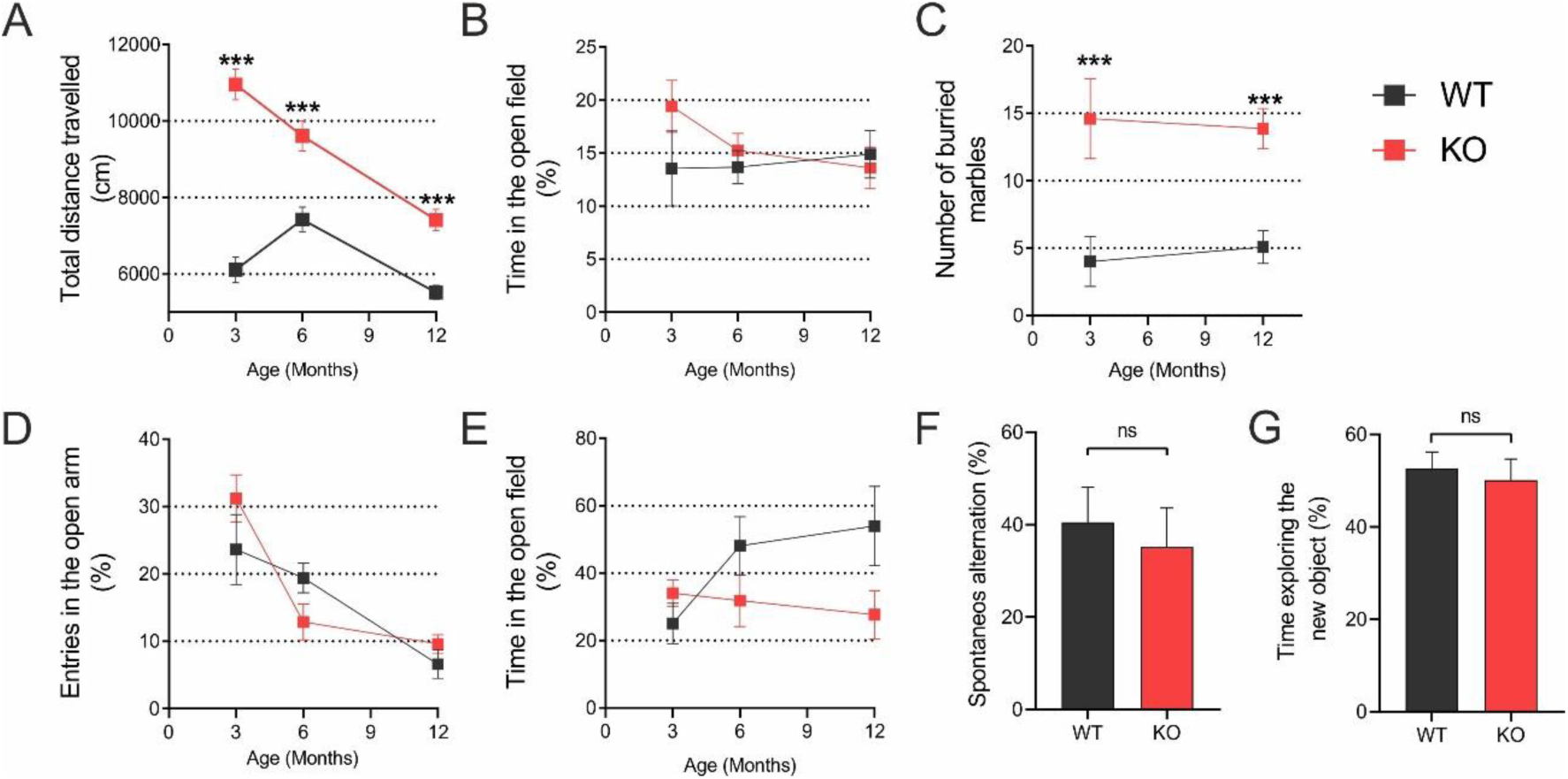
Behavioral assessment of Mrgprd deficient mice. A) Total distance traveled in the open field test (OFT) was significantly increased in MrgD-deficient mice, indicating motor hyperactivity. (B, C) No significant differences were observed in anxiety-like behavior between groups, as assessed by the percentage of entries and time spent in the open arms of the elevated plus maze. (D) MrgD-deficient mice buried significantly more marbles in the marble burying test, suggesting a compulsive-like phenotype. (E) The Y-maze test showed no differences in spontaneous alternation, indicating preserved spatial working memory. (F) In the novel object recognition (NOR) test, both genotypes showed similar discrimination between familiar and novel objects, indicating no deficits in recognition memory.

Following the analysis of motor activity and anxiety-like behavior, we assessed memory function using a series of cognitive tests. To evaluate long and short-term memory, we performed the Y-maze and novel object recognition (NOR) tests. In the Y-maze test revealed no significant changes in spontaneous alternation behavior (Figure 6E). Similarly, the NOR test, MrgD-deficient mice showed no differences in their ability to discriminate between familiar and novel objects (Figure 6F). Altogether, the behavioral tests indicate motor hyperactivity and a highly compulsive-like behavior, without deficits in short- or long-term memory (Figure 6).

## DISCUSSION

Our results demonstrate that MrgD receptor deficiency has a relevant impact on the regulation of synaptic neurotransmission, affecting multiple levels of neuronal communication. Integrative analysis of the proteome and post-translational modifications revealed significant alterations in proteins involved in the synaptic vesicle exocytosis machinery, as well as in fundamental components of dopaminergic and glutamatergic signaling. The molecular data were corroborated by functional experiments that demonstrated a reduction in the rate of vesicle exocytosis in synaptic preparations of KO mice, evidencing a direct modulatory role of the MrgD receptor in neurotransmitter release. At the behavioral level, the absence of MrgD resulted in a phenotype characterized by motor hyperactivity, increased compulsive-like behaviors and impaired olfactory memory, reinforcing the functional relevance of the receptor in the modulation of neurochemical circuits associated with motor control and cognition.

Modulation of neuronal communication is often mediated by GPCRs located in the presynaptic terminal. Among the best characterized are muscarinic, glutamatergic, opioid, cannabinoid, dopaminergic and serotonergic receptors, all recognized for their ability to regulate neurotransmitter release through multiple intracellular signaling pathways [34, 35]. These receptors act mainly via Gα subunits, modulating the activity of kinases such as PKA and PKC, which in turn promote the phosphorylation of key proteins capable of altering the efficiency of synaptic exocytosis [36]. Interestingly, kinase enrichment analysis using the differentially phosphorylated proteins in the KO mice revealed an enrichment of PKA, PKC, and CAMK2B (Figure 3B). These data provide clues to the possible mechanisms of this receptor signaling in the nigrostriatal pathway and its effects on synaptic vesicle exocytosis.

N-glycosylation is recognized as a critical modulator of synaptic transmission [37]. This post-translational modification of synaptic vesicle proteins and membrane receptors can significantly alter their functional activity independently of agonist activation, allowing fine and adaptive control of neuronal excitability and synaptic signaling [38, 39]. In addition to synaptic homeostasis, N-glycosylation plays essential roles in embryonic development and in the modulation of motor circuits [40, 41].

In this study, analysis of the N-glycoproteome revealed a significant alteration in the glycosylation of the metabotropic GABAB receptor (Gabbr2 subunit) in MrgD-KO mice (Figure 3C). This finding is particularly relevant given that modifications in the same glycosylation sites of Gabbr2 have been previously identified in murine models and in the brains of patients with Alzheimer’s disease, suggesting a possible link between MrgD receptor dysfunction, modulation of GABA-mediated inhibitory signaling and the pathophysiology of neurodegenerative disorders [42]. Additionally, alterations in the glycosylation of other key proteins involved in neurotransmission were observed, including excitatory amino acid transporter type 2 (Slc1a2) at Asn-159, syntaxin-1A (Sytx1a) at Asn-107, Stxbp1 (Munc18) at Asn-198, and Synapsin-1 (Syn1) at Asn-669 (Figure 3C). Although there is still no direct evidence in the literature associating the glycosylation of these specific sites with behavioral or pathological phenotypes, mutations in Slc1a2 are known to be implicated in neurodegenerative diseases, epilepsy, and attention deficit hyperactivity disorder (Figure S1) [43]. The presence of alterations at these sites suggests that glycosylation may impact the function of these proteins, especially considering their central role in glutamate uptake and synaptic vesicle cycling. These results point to a possible functional contribution of anomalous glycosylation in the neurobehavioral phenotype observed in animals deficient for the MrgD receptor.

The integration of proteomic, phosphoproteomic, and N-glycoproteomic data revealed a comprehensive picture of the molecular dysregulation caused by the absence of MrgD in the substantia nigra. Notably, dopaminergic, glutamatergic, and circadian neurotransmission pathways were concomitantly altered at the three levels analyzed (Figure 3C), suggesting synaptic dysfunction at multiple molecular levels with repercussions on motor tone, behavioral regulation, and physiological rhythms [44]. Changes in these two types of neurotransmission in the nigrostriatal pathway have been well characterized in the control of motor activity and compulsivity [45, 46], indicating that such molecular changes are directly associated with the phenotypes of motor hyperactivity and compulsive behavior observed in KO mice (Figure 6).

A particularly relevant finding of this study was the functional selectivity of the MrgD receptor in response to different agonists. We observed that β-alanine, at concentrations (10⁻⁴ M), did not promote significant changes in synaptic exocytosis (Figure 5A and 5B). In contrast, alamandine (10^-7^ M) significantly increased the release of synaptic vesicles in both brain regions analyzed (Figure 5A and 5B). This distinction suggests a selective pharmacological profile of MrgD, possibly dependent on multiple factors. First, differences in affinity and efficacy between the ligands may explain the divergent effects. Alamandine, an endogenous peptide of the renin-angiotensin system, may have greater affinity for MrgD or induce a more efficient activation of intracellular pathways. Furthermore, it is plausible that the two agonists promote different receptor conformations, leading to differential coupling to G proteins. Alamandine could favor signaling through Gs or Gq pathways (Figure 3B), increasing neurotransmitter release, while β-alanine could preferentially activate inhibitory (Gi/o) pathways [47], with no or suppressive impact on exocytosis (Figure 5). These findings reinforce the idea that MrgD may act as a fine modulator of synaptic function, responding selectively to endogenous stimuli with physiological relevance.

The specificity of the pro-exocytic effect of alamandine was confirmed by two complementary approaches. First, MrgD-deficient (KO) mice showed no response to alamandine treatment (Figure 5C and 5D), demonstrating that the presence of the receptor is essential for facilitating exocytosis. Second, administration of the partial antagonist DPRO (10^-6^ M) partially blocked the effect of alamandine (Figure 5E and 5F), strengthening the evidence that the effect is mediated directly by MrgD activation.

The behavioral data reinforce the functional relevance of the molecular and physiological findings presented in this study. Mice deficient for the MrgD receptor exhibited pronounced motor hyperactivity and compulsive behavior phenotypes (Figure 6), characteristics compatible with dysfunctions in dopaminergic and glutamatergic neurotransmission in the nigrostriatal pathway, as previously described in the literature [46, 48]. This failure in dopaminergic neurotransmission appears to explain the motor impairment observed in KO animals, leading to functional impairments in this pathway critical for the control of motor activity [49]. Behavioral analysis performed over 12 months revealed two distinct patterns of change: persistent compulsive behavior that remained stable with aging, and motor hyperactivity that progressively decreased with age. These findings indicate that the absence of MrgD is associated with long-lasting neurobehavioral changes, with possible relevance for the understanding of neuropsychiatric and neurodegenerative disorders.

The chronic compulsivity observed in KO mice may be related to dysfunctions in the corticostriatal circuits, responsible for the control of repetitive behaviors and the inhibition of actions [50, 51]. Considering that the striatum is a key region for the control of stereotyped actions [52] and that the data demonstrated impairment in synaptic exocytosis in this area, it is plausible that the alteration in dopaminergic and glutamatergic neurotransmission favors the development of these compulsive behaviors. Furthermore, there is evidence that this type of behavior may emerge spontaneously in the early stages of nigrostriatal degeneration [53].

On the other hand, motor hyperactivity showed a gradual decline with aging, a pattern consistent with murine models that initially exhibit hyperlocomotion and, over time, develop motor deficits associated with loss of dopaminergic function [54]. This behavior may reflect a progressive reduction in the plasticity of the nigrostriatal pathway in the absence of MrgD, contributing to the decrease in spontaneous motor activity throughout life. This pattern is particularly relevant for understanding the early mechanisms that precede the development of diseases such as Parkinson’s, suggesting that the MrgD receptor may play an essential modulatory role in the maintenance of neurotransmission and motor function throughout aging.

Thus, the findings of this study reveal a new and significant function for the MrgD receptor in the central nervous system, expanding its role beyond the previously established peripheral nociception [4]. By demonstrating that activation or absence of MrgD directly affects synaptic exocytosis and translates into behavioral changes, such as compulsivity and motor hyperactivity, these results establish a clear connection between the molecular signaling mediated by this receptor and complex neurobehavioral manifestations. Thus, MrgD is consolidated as a key modulator of neurotransmission in the nigrostriatal pathway, positioning itself as a promising target for the development of new therapeutic approaches aimed at the treatment of neuropsychiatric disorders characterized by dysfunctions in motor and behavioral control.

### Limitations of the Study

Some limitations of this study should be considered. First, although the multiomics approach allowed a comprehensive view of the molecular regulation associated with the MrgD receptor, the analyses were performed on whole tissue from the substantia nigra and striatum, which may mask specific alterations in distinct cell types. Future approaches with cell separation, such as single-cell proteomics and/or spatial proteomics, could offer greater spatial and functional resolution. Furthermore, the functional interpretation of post-translational modification data, especially N-glycosylation, is limited by the current knowledge in literature about the modified sites. Although we identified consistent alterations in key neurotransmission proteins, the specific functionality of these glycosylation events is not yet fully elucidated and requires experimental validation by targeted functional assays. Finally, the study focused mainly on murine models, which may limit the direct extrapolation of the results to human clinical settings. Complementary investigations with human samples or patient-derived cellular models could contribute to validate the translational relevance of the findings.

---

**Figure S1.**
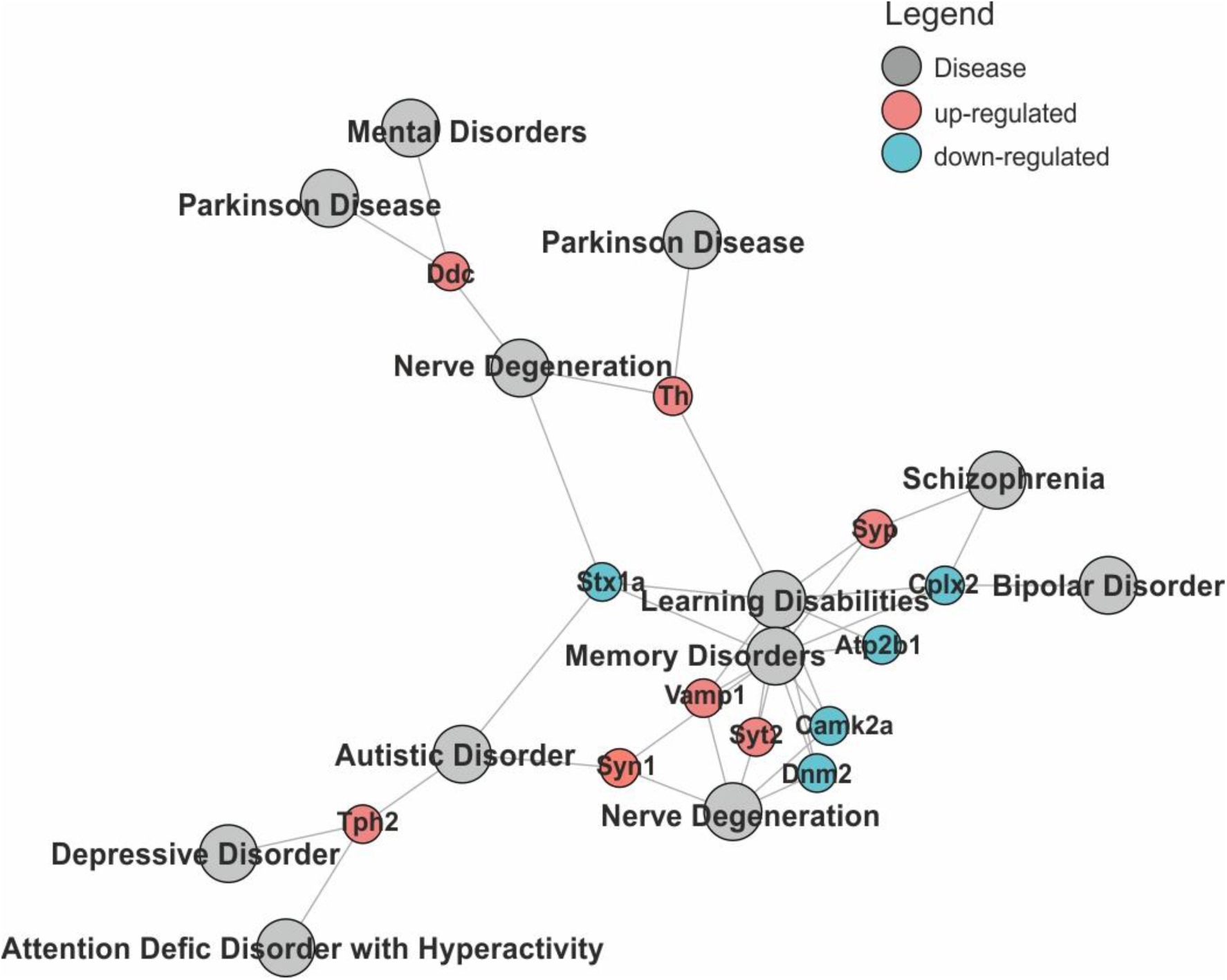
Disease enrichment using regulated proteins in the SN. The enrichment was performed using the Comparative Toxicogenomics Database (CTD). The graph was built using “igraph” package in R environment.

## References

1. Lan L, Xu M, Li J, et al (2020) Mas-related G protein-coupled receptor D participates in inflammatory pain by promoting NF-κB activation through interaction with TAK1 and IKK complex. Cell Signal 76:109813. 10.1016/j.cellsig.2020.109813

2. Guo C, Jiang H, Huang C-C, et al (2023) Pain and itch coding mechanisms of polymodal sensory neurons. Cell Rep 42:113316. 10.1016/j.celrep.2023.113316

3. Wang L, Su X, Yan J, et al (2023) Involvement of Mrgprd-expressing nociceptors-recruited spinal mechanisms in nerve injury-induced mechanical allodynia. iScience 26:106764. 10.1016/j.isci.2023.106764

4. Liu Q, Sikand P, Ma C, et al (2012) Mechanisms of itch evoked by β-alanine. J Neurosci Off J Soc Neurosci 32:14532–14537. 10.1523/JNEUROSCI.3509-12.2012

5. Lautner RQ, Villela DC, Fraga-Silva RA, et al (2013) Discovery and characterization of alamandine: a novel component of the renin-angiotensin system. Circ Res 112:1104–1111. 10.1161/CIRCRESAHA.113.301077

6. Almeida-Santos AF, de Melo LA, Gonçalves SCA, et al (2021) Alamandine through MrgD receptor induces antidepressant-like effect in transgenic rats with low brain angiotensinogen. Horm Behav 127:104880. 10.1016/j.yhbeh.2020.104880

7. Valenzuela R, Rodriguez-Perez AI, Costa-Besada MA, et al (2021) An ACE2/Mas-related receptor MrgE axis in dopaminergic neuron mitochondria. Redox Biol 46:102078. 10.1016/j.redox.2021.102078

8. Hami J, von Bohlen Und Halbach V, Tetzner A, et al (2021) Localization and expression of the Mas-related G-protein coupled receptor member D (MrgD) in the mouse brain. Heliyon 7:e08440. 10.1016/j.heliyon.2021.e08440

9. Lanciego JL, Luquin N, Obeso JA (2012) Functional Neuroanatomy of the Basal Ganglia. Cold Spring Harb Perspect Med 2:a009621. 10.1101/cshperspect.a009621

10. Zou L, Tian Y, Zhang Z (2021) Dysfunction of Synaptic Vesicle Endocytosis in Parkinson’s Disease. Front Integr Neurosci 15:619160. 10.3389/fnint.2021.619160

11. Özçete ÖD, Banerjee A, Kaeser PS (2024) Mechanisms of neuromodulatory volume transmission. Mol Psychiatry 29:3680–3693. 10.1038/s41380-024-02608-3

12. Sauvola CW, Littleton JT (2021) SNARE Regulatory Proteins in Synaptic Vesicle Fusion and Recycling. Front Mol Neurosci 14:733138. 10.3389/fnmol.2021.733138

13. Channer B, Matt SM, Nickoloff-Bybel EA, et al (2023) Dopamine, Immunity, and Disease. Pharmacol Rev 75:62–158. 10.1124/pharmrev.122.000618

14. Teleanu RI, Niculescu A-G, Roza E, et al (2022) Neurotransmitters-Key Factors in Neurological and Neurodegenerative Disorders of the Central Nervous System. Int J Mol Sci 23:5954. 10.3390/ijms23115954

15. Meier F, Brunner A-D, Koch S, et al (2018) Online Parallel Accumulation-Serial Fragmentation (PASEF) with a Novel Trapped Ion Mobility Mass Spectrometer. Mol Cell Proteomics MCP 17:2534–2545. 10.1074/mcp.TIR118.000900

16. Engholm-Keller K, Larsen MR (2013) Technologies and challenges in large-scale phosphoproteomics. Proteomics 13:910–931. 10.1002/pmic.201200484

17. Betz WJ, Bewick GS (1993) Optical monitoring of transmitter release and synaptic vesicle recycling at the frog neuromuscular junction. J Physiol 460:287–309. 10.1113/jphysiol.1993.sp019472

18. Harata N, Ryan TA, Smith SJ, et al (2001) Visualizing recycling synaptic vesicles in hippocampal neurons by FM 1-43 photoconversion. Proc Natl Acad Sci U S A 98:12748– 12753. 10.1073/pnas.171442798

19. Kay AR, Alfonso A, Alford S, et al (1999) Imaging synaptic activity in intact brain and slices with FM1-43 in C. elegans, lamprey, and rat. Neuron 24:809–817. 10.1016/s0896-6273(00)81029-6

20. De Benedetto GE, Fico D, Pennetta A, et al (2014) A rapid and simple method for the determination of 3,4-dihydroxyphenylacetic acid, norepinephrine, dopamine, and serotonin in mouse brain homogenate by HPLC with fluorimetric detection. J Pharm Biomed Anal 98:266–270. 10.1016/j.jpba.2014.05.039

21. Kraeuter A-K, Guest PC, Sarnyai Z (2019) The Open Field Test for Measuring Locomotor Activity and Anxiety-Like Behavior. Methods Mol Biol Clifton NJ 1916:99–103. 10.1007/978-1-4939-8994-2_9

22. Komada M, Takao K, Miyakawa T (2008) Elevated plus maze for mice. J Vis Exp JoVE 1088. 10.3791/1088

23. Kraeuter A-K, Guest PC, Sarnyai Z (2019) The Y-Maze for Assessment of Spatial Working and Reference Memory in Mice. Methods Mol Biol Clifton NJ 1916:105–111. 10.1007/978-1-4939-8994-2_10

24. Leger M, Quiedeville A, Bouet V, et al (2013) Object recognition test in mice. Nat Protoc 8:2531–2537. 10.1038/nprot.2013.155

25. Courtney NA, Bao H, Briguglio JS, Chapman ER (2019) Synaptotagmin 1 clamps synaptic vesicle fusion in mammalian neurons independent of complexin. Nat Commun 10:4076. 10.1038/s41467-019-12015-w

26. Vardar G, Chang S, Arancillo M, et al (2016) Distinct Functions of Syntaxin-1 in Neuronal Maintenance, Synaptic Vesicle Docking, and Fusion in Mouse Neurons. J Neurosci Off J Soc Neurosci 36:7911–7924. 10.1523/JNEUROSCI.1314-16.2016

27. Makke M, Pastor-Ruiz A, Yarzagaray A, et al Key determinants of the dual clamp/activator function of Complexin. eLife 12:RP92438. 10.7554/eLife.92438

28. Ge Z, Gu Y, Han Q, et al (2016) Targeting High Dynamin-2 (DNM2) Expression by Restoring Ikaros Function in Acute Lymphoblastic Leukemia. Sci Rep 6:38004. 10.1038/srep38004

29. Daubner SC, Le T, Wang S (2011) Tyrosine hydroxylase and regulation of dopamine synthesis. Arch Biochem Biophys 508:1–12. 10.1016/j.abb.2010.12.017

30. Bertoldi M (2014) Mammalian Dopa decarboxylase: structure, catalytic activity and inhibition. Arch Biochem Biophys 546:1–7. 10.1016/j.abb.2013.12.020

31. Kulikova EA, Kulikov AV (2019) Tryptophan hydroxylase 2 as a therapeutic target for psychiatric disorders: focus on animal models. Expert Opin Ther Targets 23:655–667. 10.1080/14728222.2019.1634691

32. de Beer AD, Legoabe LJ, Petzer A, Petzer JP (2021) The inhibition of catechol O-methyltransferase and monoamine oxidase by tetralone and indanone derivatives substituted with the nitrocatechol moiety. Bioorganic Chem 114:105130. 10.1016/j.bioorg.2021.105130

33. Ghirardi M, Braha O, Hochner B, et al (1992) Roles of PKA and PKC in facilitation of evoked and spontaneous transmitter release at depressed and nondepressed synapses in Aplysia sensory neurons. Neuron 9:479–489. 10.1016/0896-6273(92)90185-g

34. Yim YY, Zurawski Z, Hamm H (2018) GPCR regulation of secretion. Pharmacol Ther 192:124–140. 10.1016/j.pharmthera.2018.07.005

35. Lovinger DM, Mateo Y, Johnson KA, et al (2022) Local modulation by presynaptic receptors controls neuronal communication and behaviour. Nat Rev Neurosci 23:191–203. 10.1038/s41583-022-00561-0

36. Epac2 Mediates cAMP-Dependent Potentiation of Neurotransmission in the Hippocampus - PubMed. https://pubmed.ncbi.nlm.nih.gov/25904804/. Accessed 9 Jul 2025

37. Scott H, Panin VM (2014) N-glycosylation in regulation of the nervous system. Adv Neurobiol 9:367–394. 10.1007/978-1-4939-1154-7_17

38. Gurevicius K, Gureviciene I, Sivukhina E, et al (2007) Increased hippocampal and cortical beta oscillations in mice deficient for the HNK-1 sulfotransferase. Mol Cell Neurosci 34:189–198. 10.1016/j.mcn.2006.10.014

39. Conroy LR, Hawkinson TR, Young LEA, et al (2021) Emerging roles of N-linked glycosylation in brain physiology and disorders. Trends Endocrinol Metab TEM 32:980–993. 10.1016/j.tem.2021.09.006

40. Ye Z, Marth JD (2004) N-glycan branching requirement in neuronal and postnatal viability. Glycobiology 14:547–558. 10.1093/glycob/cwh069

41. Pradeep P, Kang H, Lee B (2023) Glycosylation and behavioral symptoms in neurological disorders. Transl Psychiatry 13:154. 10.1038/s41398-023-02446-x

42. Zhang Q, Ma C, Chin L-S, Li L (2020) Integrative glycoproteomics reveals protein N-glycosylation aberrations and glycoproteomic network alterations in Alzheimer’s disease. Sci Adv 6:eabc5802. 10.1126/sciadv.abc5802

43. Todd AC, Hardingham GE (2020) The Regulation of Astrocytic Glutamate Transporters in Health and Neurodegenerative Diseases. Int J Mol Sci 21:9607. 10.3390/ijms21249607

44. David HN (2009) Towards a reconceptualization of striatal interactions between glutamatergic and dopaminergic neurotransmission and their contribution to the production of movements. Curr Neuropharmacol 7:132–141. 10.2174/157015909788848893

45. Karthik S, Sharma LP, Narayanaswamy JC (2020) Investigating the Role of Glutamate in Obsessive-Compulsive Disorder: Current Perspectives. Neuropsychiatr Dis Treat 16:1003– 1013. 10.2147/NDT.S211703

46. Pittenger C, Bloch MH, Williams K (2011) Glutamate abnormalities in obsessive compulsive disorder: neurobiology, pathophysiology, and treatment. Pharmacol Ther 132:314–332. 10.1016/j.pharmthera.2011.09.006

47. Suzuki S, Iida M, Hiroaki Y, et al (2022) Structural insight into the activation mechanism of MrgD with heterotrimeric Gi-protein revealed by cryo-EM. Commun Biol 5:1–13. 10.1038/s42003-022-03668-3

48. Andersson DR, Björnsson E, Bergquist F, Nissbrandt H (2010) Motor activity-induced dopamine release in the substantia nigra is regulated by muscarinic receptors. Exp Neurol 221:251–259. 10.1016/j.expneurol.2009.11.011

49. Masuo Y, Morita M, Oka S, Ishido M (2004) Motor hyperactivity caused by a deficit in dopaminergic neurons and the effects of endocrine disruptors: a study inspired by the physiological roles of PACAP in the brain. Regul Pept 123:225–234. 10.1016/j.regpep.2004.05.010

50. Vicente AM, Martins GJ, Costa RM (2020) Cortico-basal ganglia circuits underlying dysfunctional control of motor behaviors in neuropsychiatric disorders. Curr Opin Genet Dev 65:151–159. 10.1016/j.gde.2020.05.042

51. Maia TV, Cooney RE, Peterson BS (2008) The neural bases of obsessive-compulsive disorder in children and adults. Dev Psychopathol 20:1251–1283. 10.1017/S0954579408000606

52. Graybiel AM, Grafton ST (2015) The striatum: where skills and habits meet. Cold Spring Harb Perspect Biol 7:a021691. 10.1101/cshperspect.a021691

53. Averbeck BB, O’Sullivan SS, Djamshidian A (2014) Impulsive and compulsive behaviors in Parkinson’s disease. Annu Rev Clin Psychol 10:553–580. 10.1146/annurev-clinpsy-032813-153705

54. Ferguson SA, Law CD, Sarkar S (2015) Chronic MPTP treatment produces hyperactivity in male mice which is not alleviated by concurrent trehalose treatment. Behav Brain Res 292:68–78. 10.1016/j.bbr.2015.05.057

