## Supplementary figures and images for "MrgD Receptor Modulates Neurotransmission in the Nigrostriatal Pathway"

### Figure S1

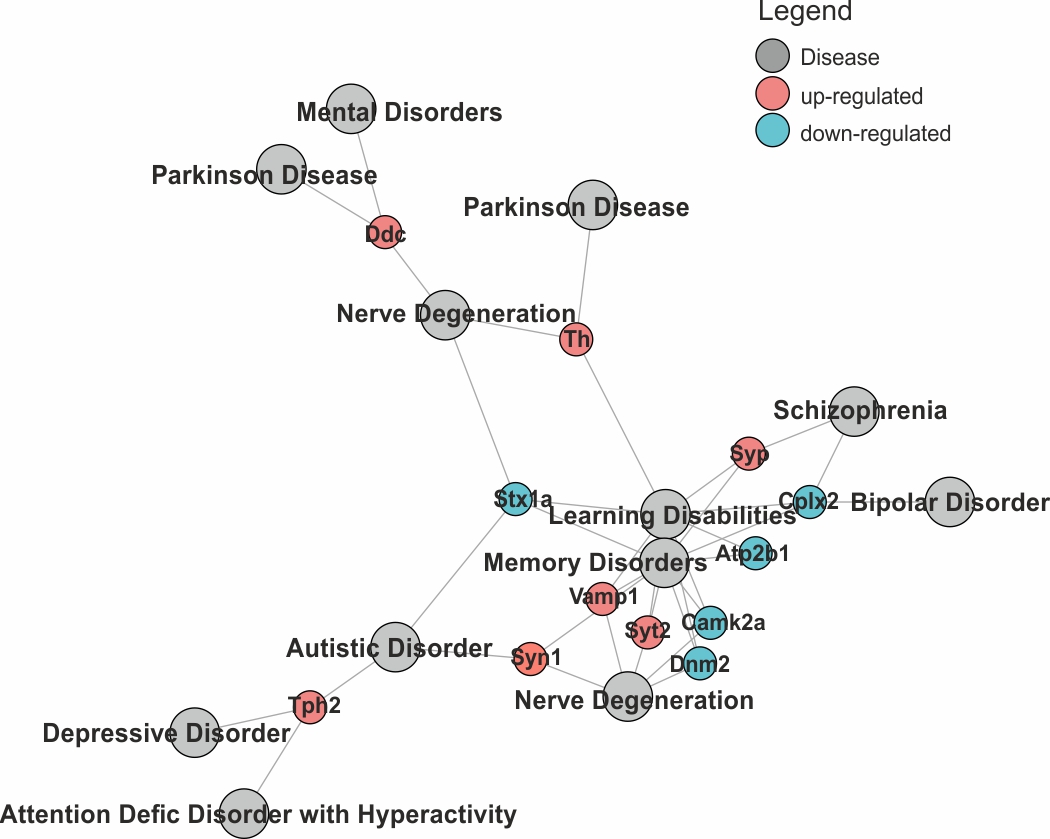
